# Measuring sub-nanometer fluctuations at microsecond temporal resolution with metal-and graphene-induced energy transfer spectroscopy

**DOI:** 10.1101/2023.05.16.540910

**Authors:** Tao Chen, Narain Karedla, Jörg Enderlein

## Abstract

Out-of-plane fluctuations, also known as stochastic displacements, of biological membranes play a crucial role in regulating many essential life processes within cells and organelles. Despite the availability of various methods for quantifying membrane dynamics, accurately quantifying complex membrane systems with rapid and tiny fluctuations, such as mitochondria, remains a challenge. In this work, we present a novel methodology that combines metal/graphene-induced energy transfer (MIET/GIET) with fluorescence correlation spectroscopy (FCS) to quantify out-of-plane fluctuations of membranes with simultaneous spatiotemporal resolution of approximately one nanometer and one microsecond that is unprecedented.

To validate the technique and spatiotemporal resolution, we measured bending undulations of model membranes. Furthermore, we demonstrate the versatility and applicability of MIET/GIET-FCS for studying diverse membrane systems, including the widely studied fluctuating membrane system of human red blood cells, as well as two unexplored membrane systems with tiny fluctuations, a pore-spanning membrane, and mitochondrial inner/outer membranes.

## 1 Introduction

Cellular membranes play a crucial role in many fundamental physiological processes, such as cell adhesion, migration, signaling, and trafficking. From a physics perspective, membranes are selectively permeable, viscoelastic interfaces that surround cells and cellular organelles. They can be thought of as two-dimensional fluids composed mostly of lipids and membrane proteins that undergo Brownian motion within the plane of the membrane. Despite their complex structures, cellular membranes behave as if they are nearly perfect elastic shells, exhibiting linear viscoelastic responses and thermally induced spatial fluctuations, which are primarily bending fluctuations known as undulations.

Researchers have only shown in the past three decades that the non-equilibrium nature of membrane fluctuations can be attributed to active sources. These sources include cytoskeletal dynamics, protein and ion pumps, lipid transport systems, ATP-driven membrane remodeling, and the membrane-fission and fusion of cargo vesicles [1–8]. Brochard and Lennon presented the first quantitative description of the flickering dynamics of red blood cell membranes. Since then, this description has been further extended in numerous works to include active force generation from point sources such as ion channels, transporters, and other factors [9–12]. However, these fluctuations have a relatively wide temporal range, and current theoretical models lack the necessary complexity to describe them quantitatively (see the review by Turlier *et al.* [13]).

Measuring cell membrane fluctuations provides valuable insights into various aspects of cellular health, including mechanical properties, disease states, drug responses, and fundamental biological processes. By comprehending these fluctuations, one can evaluate cellular function, detect biomarkers for diseases, optimize drug development and screening, and enhance our understanding of basic biological processes. This knowledge will have far-reaching implications for fields such as cell biology, biophysics, biomedical research, and medicine, ultimately resulting in improved diagnostics, treatments, and overall human health.

Several methods have been used to measure membrane fluctuations [14]. Camera-based wide-field detection methods such as contour analysis [15], fluorescence interference contrast (FLIC) microscopy [16–18], flicker spectroscopy [19], diffraction phase microscopy [3], and reflection interference contrast microscopy (RICM) [20–22] capture membrane dynamics with millisecond temporal resolution (normally around ∼ 10 ms), and with approximately 10 nm resolution in the case of RICM. In contrast, single point detection methods such as time-resolved membrane fluctuation spectroscopy (TRMFS) [4, 23] or dynamic optical displacement spectroscopy (DODS) [14] achieve temporal resolutions of ∼ 1 − 10 µs, but do not provide the spatial information of camera-based methods.

Most experimental work so far has focused on measuring membrane fluctuations on deflated giant unilamellar vesicles (GUVs) or red blood cells under stringent physiological conditions that lead to large amplitude bending fluctuations between 50 nm to 100 nm due to vanishing membrane surface tension. However, membranes of adherent cells or organelles that are under tension show tiny fluctuations with an amplitude below 10 nm at frequencies above 10 kHz [24]. However, current state-of-the-art techniques do not achieve the necessary spatial and temporal resolution to capture such fluctuations.

In this study, we present a novel method for quantifying nanoscale and sub-nanoscale membrane fluctuations with sub-millisecond temporal resolution using metal-/graphene-induced energy transfer (MIET/GIET). MIET/GIET relies on the electrodynamic coupling of a fluorescent emitter to either surface plasmons in a thin metal film (MIET) [25, 26] or excitons in a single sheet of graphene (GIET) [27, 28], see Figures 1c,d. This coupling leads to a strong modulation of the excited state lifetime of the fluorescent dye that depends on its distance from the metal or graphene layers. By measuring the excited state lifetime of the dye, we can determine its distance from the metal or graphene layers, and convert this lifetime into a distance using the well-understood physics of electrodynamics. This method is a significant improvement over previous techniques and provides unprecedented insight into the dynamics of cellular membranes.

**Fig. 1.**
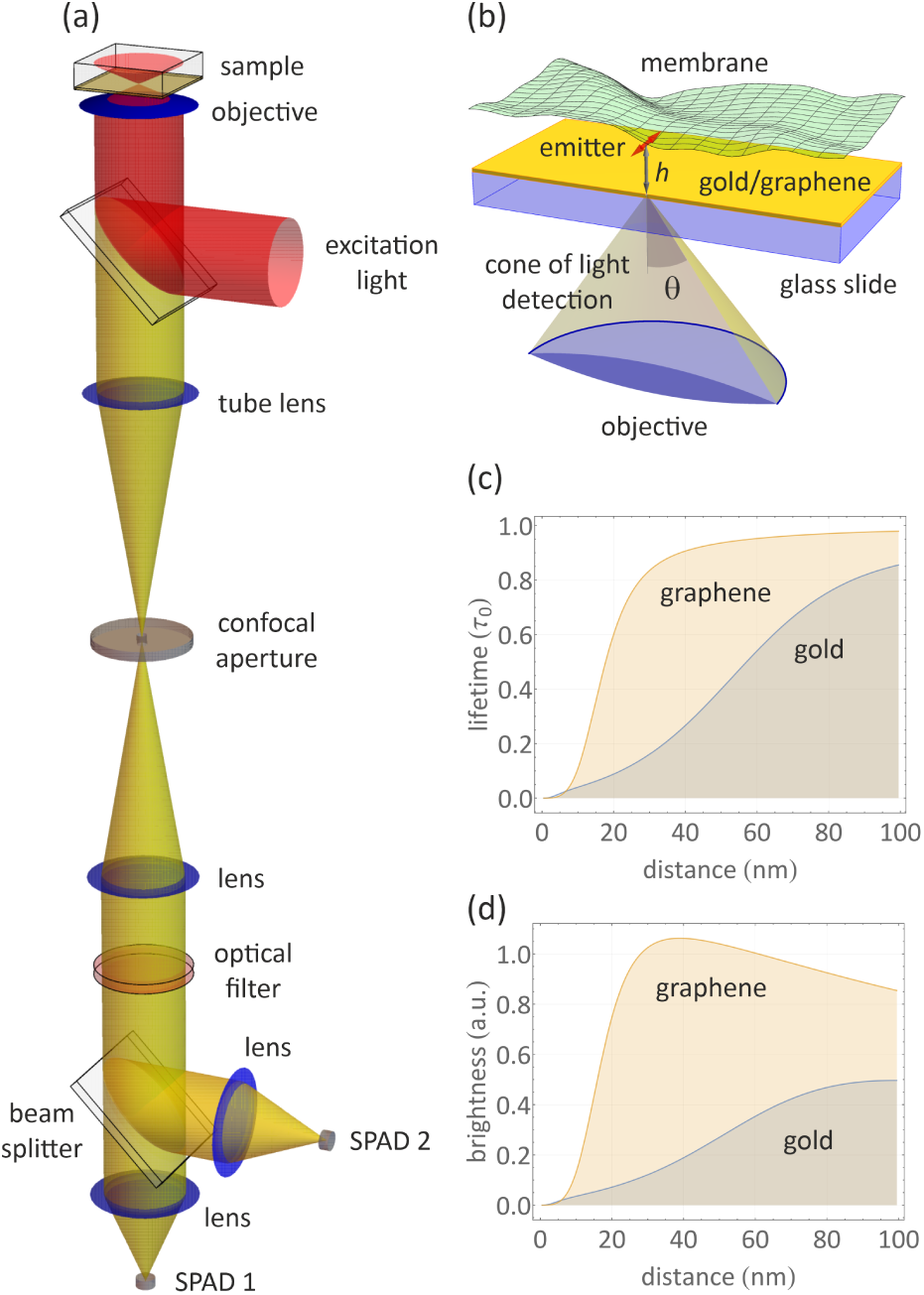
(a) Schematic of the MIET/GIET measurement setup. Fluorescence excitation and detection are performed through the coverslip from below, using a high numerical aperture objective. A conventional confocal fluorescence microscope equipped with a multichannel picosecond event timer for measuring fluorescence lifetimes is used for the fluorescence intensity and lifetime measurements. (b) Schematic of a fluctuating membrane above a gold film. (c) Calculated relative fluorescence lifetime *τ/τ*_0_ of an emitter as a function of its distance *h* from the surface of a 10 nm gold film or single sheet of graphene on glass. (d) Calculated relative brightness of an emitter as a function of its distance *h* from the surface of a 10 nm gold film or single sheet of graphene on glass. All calculations were performed for an emitter with maximum emission wavelength of 680 nm, fluorescence quantum yield of 0.8, and random orientation.

In previous studies, MIET has been primarily used for structural analysis, such as mapping the topography of the basal membrane of living cells [26], reconstructing focal adhesions and stress fibers in three dimensions [29], measuring the distance between the inner and outer envelope of the nucleus [30], visualizing the dynamics of epithelial-mesenchymal transitions (EMT) [31], and enabling single-molecule localization and co-localization along the optical axis [32, 33]. Additionally, MIET has been utilized to map the basal membrane and lamellipodia of human aortic endothelial cells [34, 35] and to achieve three-dimensional isotropic resolution imaging of microtubules and clathrin pits in combination with SMLM [36]. By replacing the thin metal film with a single sheet of graphene (Graphene-Induced Energy Transfer or GIET), the localization accuracy can be improved by an order of magnitude within ∼25 nm from the substrate [27]. GIET has been employed to measure the thickness of supported lipid bilayers with different lipid compositions, enabling an accuracy of a few°angstroms [28], as well as to observe small changes in mitochondrial membrane organization upon respiration [37], or to quantify cholesterol-induced thickness changes of lipid bilayers [38].

To measure rapid membrane fluctuations, we combine MIET/GIET with fluorescence correlation spectroscopy (FCS) [39], using the setup shown in Figure 1a. Our method is based on the fact that MIET/GIET not only modulates the fluorescence lifetime of dyes close to the substrate (Figure 1c), but also the brightness (Figure 1d), leading to strong fluctuations in fluorescence intensity when a fluorescently labeled membrane close to a MIET or GIET substrate fluctuates perpendicular to the substrate’s surface, see Figure 1b. FCS allows for the precise measurement and quantification of these fluctuations, which appear as a partial correlation decay on the time-scale of the membrane fluctuations. This approach is particularly useful for studying rapid membrane dynamics with sub-millisecond temporal resolution and nanoscale and sub-nanoscale spatial accuracy.

We demonstrate the method and validate its spatial and temporal resolution by measuring membrane fluctuations of model giant unilamellar vesicles (GUVs). We then apply MIET/GIET-FCS to three different systems: (i) investigating the flickering of red blood cell (RBC) membranes with and without ATP; (ii) studying the fluctuations of pore-spanning membranes with small fluctuation amplitudes at high frequency; and (iii) exploring the undulations of the outer and inner membranes in active and resting-state mitochondria.

## 2 Theory

Consider an electric dipole emitter that fluoresces (emits light) and is positioned at a distance *h* above the surface of a MIET/GIET substrate, as depicted in Figure 1b. When the distance *h* decreases to a certain value (below 200 nm for a gold film, or below 25 nm for graphene), the electric near field of the emitter can stimulate surface plasmons in the metal film or excitons in graphene, causing the emitter to de-excite more quickly from its excited state to the ground state. This phenomenon can be studied by solving Maxwell’s equations for the electromagnetic field of the electric dipole emitter in the presence of the MIET/GIET substrate, as outlined in Refs. [28, 32]. This computation yields several important quantities.

Firstly, the total emission power *S*(*h, β*) of the emitter can be determined as a function of its distance *h* from the substrate and its orientation, described by the angle *β* between its dipole axis and the vertical direction. This is achieved by integrating the (time-averaged) Poynting vector, which is the cross product of the electric and magnetic field vectors, multiplied by the speed of light and divided by 8*π*, across two horizontal planes within the emitter’s medium (buffer solution) that enclose the emitter from both sides.

Secondly, the total far-field emission *N* (*h, β*) that is radiated into the lower half space (glass over slide) can be calculated by integrating the Poynting vector over the interface between the glass and the metal/graphene layer.

Lastly, the relative proportion *C*(*h, β*) of the emission that is collected by the objective lens, which is the part that is radiated into its cone of light detection, can be determined by integrating the angular distribution of emission (the Poynting vector as a function of the emission angle) over this cone of light detection.

This emission power *S*(*h, β*) is proportional to the *radiative* transition rate of a fluorescent molecule from its excited to its ground state. By knowing the quantum yield of fluorescence *ϕ* (ratio of radiative to total transition rate), one can calculate the observable fluorescence lifetime *τ_f_* as follows:

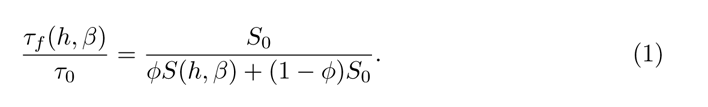

In this equation, *S*_0_ = *cnk*^4^*p*^2^*/*3 represents the emission power of an electric dipole emitter with unity quantum yield in a homogeneous unbounded dielectric medium with refractive index *n* (which is equal to the refractive index of the medium above the MIET substrate), where *c* is the speed of light, *k*_0_ is the wave vector in vacuum, and *p* is the amplitude of the dipole moment vector. *τ*_0_ is the corresponding freespace fluorescence lifetime. The angular dependence of *S*(*h, β*) can be decomposed as *S*(*h, β*) = *S_⊥_*(*h*) cos^2^ *β* + *S_║_*(*h*) sin^2^ *β*, where *S_⊥_*(*h*) and *S_∥≑_*(*h*) are the radiative emission rates of emitters oriented perpendicular and parallel to the substrate, respectively, and are now functions of *h* only. When a molecule is freely rotating (i.e., its reorientation rate is much faster than its fluorescence lifetime), the full emission rate can be simplified to *S*(*h*) = [*S_⊥_*(*h*) + 2*S_║_*(*h*)]*/*3 after averaging over all possible orientations.

The curves in Figure 1c demonstrate the calculation of the relative lifetime *τ_f_* (*h*)*/τ*_0_ for a rapidly rotating dye with a quantum yield of *ϕ* = 0.8 emitting at *λ*_em_ = 680,nm as a function of height *h* over a gold-coated and a graphene substrate. As can be observed, the fluorescence lifetime increases as a monotonic function within a distance of up to approximately 150 nm for the gold layer and up to 25 nm for graphene.

In nearly all applications of MIET and GIET thus far, the dependency of lifetime on distance has been utilized to localize an emitter along the optical axis by measuring its lifetime. However, the near-field coupling and energy transfer of an emitter to a metal or graphene layer not only modulates its excited-state lifetime but also affects its observable brightness. By determining the total far-field emission *N* (*h, β*) into the glass and its relative portion *C*(*h, β*) collected by the objective, one can calculate the brightness value *b*(*h*) of a randomly oriented or rapidly rotating emitter using the following equation (up to some proportionality constant):

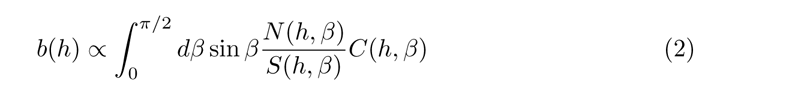

Similar to the case of *S*(*h, β*), the function *N* (*h, β*) can be decomposed as *N_⊥_*(*h, β*) cos^2^ *β* + *N_║_*(*h, β*) sin^2^ *β*, and similarly for *C*(*h, β*). A comprehensive description of all the mathematical details of this calculation is presented in Ref. [40].

The graph in Figure 1d illustrates the relative brightness *b*(*h*) of a dipole under the same conditions as the lifetime calculations, but with an objective having an N.A. of 1.49 and *n*_imm_ of 1.52. It should be noted that when the dipole is far from the metal or graphene surface and the energy transfer is negligible, the relative brightness approaches the net transparency of the substrate. We verified the lifetime and relative brightness calculations using additional experiments where we measured these parameters on bilayers for various thickness of silica spacers (see Supplementary Figure 1 and Supplementary Note 1).

The distance-dependent detectable fluorescence brightness is of particular significance in the current study, as it enables the combination with FCS to measure minute vertical membrane fluctuations near a MIET/GIET substrate in a fast and precise manner. In the absence of a MIET/GIET substrate, FCS would primarily capture intensity fluctuations resulting from the lateral diffusion of labeled lipids through the confocal volume. However, in the presence of a MIET/GIET substrate, these intensity fluctuations are also influenced by vertical position fluctuations of the emitters. Furthermore, high label densities of fluorescently labeled lipid membranes render the impact of lateral molecular diffusion on the observable fluorescence intensity completely negligible, and the resulting auto-correlation function (ACF) measured in FCS experiments is solely determined by the vertical position fluctuations of the membrane. The intensity correlation function *g_I_* (*t*) can be expressed as the time average of the product of intensities *I*(*t^′^*) and (*t^′^* + *t*), denoted by ⟨*δI*(*t^′^*)*δI*(*t^′^* + *t*)⟩*_t_′* where *δI*(*t*) is the difference between the intensity *I*(*t*) and its time-averaged value, and the angular brackets with index *t^′^* indicate the time average. When the time-dependent height of the membrane, *h*(*t*) = *h*_0_ +*δh*(*t*), falls within a range where the intensity-versus-height relationship can be approximated by a linear function, the height correlation function

*gh*(*t*) = ⟨*δh*(*t^′^*)*δh*(*t^′^* + *t*)⟩*_t_′* can be related to *g_I_* (*t*) through the approximation

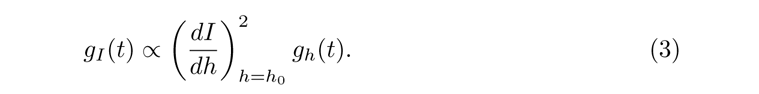

Thus, measuring the intensity correlation function *g_I_* does, in the linear regime, allow for determining the height correlation function *g_h_*. The average height *h*_0_ itself can be estimated from the average fluorescence lifetime using the lifetime-versus-height MIET/GIET relationship, see Figure 1c. As can be seen from Figure 1d, the distance range where the r.h.s. of eq. (3) is an acceptable approximation is ca. 30-70 nm for MIET (gold layer), and ca. 10-25 nm for GIET (graphene). We also used GUV membrane measurements for experimentally checking the validity of assuming a linear relationship between height and intensity, see Supplementary Figure 2.

The ordinate of the height correlation func<u>tion</u> at lag time *t* = 0 yields the root mean square amplitude of displacement *ψ* = ⟨*δh*^2^⟩ and the relaxation time *τ^∗^* where this function has fallen of to one half of its maximum value.

## 3 Results

### 3.1 MIET-FCS

In the next two subsections, we will use MIET for measuring membrane fluctuations of GUVs and cells. In all these experiment, the used MIET substrate has always the same structure: On a glass cover slide (refractive index 1.52), we deposited a 10 nm thick gold film which was covered by a 10 nm thick protecting quartz layer.

#### 3.1.1 Membrane fluctuations in GUVs

We assessed the performance of MIET-FCS by measuring membrane fluctuations of GUVs labeled with fluorescent markers [14]. A typical geometry of the experiment is shown in Figure 2a. We first showed that a densely labeled membrane leads to a flat ACF with no lag-time dependence. For this purpose, we prepared GUVs from SOPC and fluorescently labeled DPPE (DPPE-Atto655) by electroformation [28] in a sucrose solution of 230 mOs/mL. We used two different concentrations of fluorescently labeled lipids (DPPE-Atto655): 10^-3^ mol% and 1 mol%. We immersed GUVs into a measurement chamber containing an isotonic solution, which had an osmolarity equivalent to that of the GUVs. Most of the GUVs settle down at the bottom of the chambers within 15 min forming a flattened contact area with the surface (see Figures 2b,c). FCS measurements were recorded at the bottom of the chamber (glass cover slide covered with BSA) with the excitation focus of the confocal microscope placed within the contact area (glass surface). Figure 2d shows a comparison of the measured intensity correlation functions for both label concentrations. At 10^-3^ mol% labeling concentration, we observed a temporally decaying correlation due to the diffusion of labeled lipids in and out of the focus, with a diffusion coefficient of 7.3 ± 0.3 µm^2^s^-1^, which is close to reported literature values [41]. At 1 mol% labeling concentration, this diffusion-related decay of the correlation function is negligible.

**Fig. 2.**
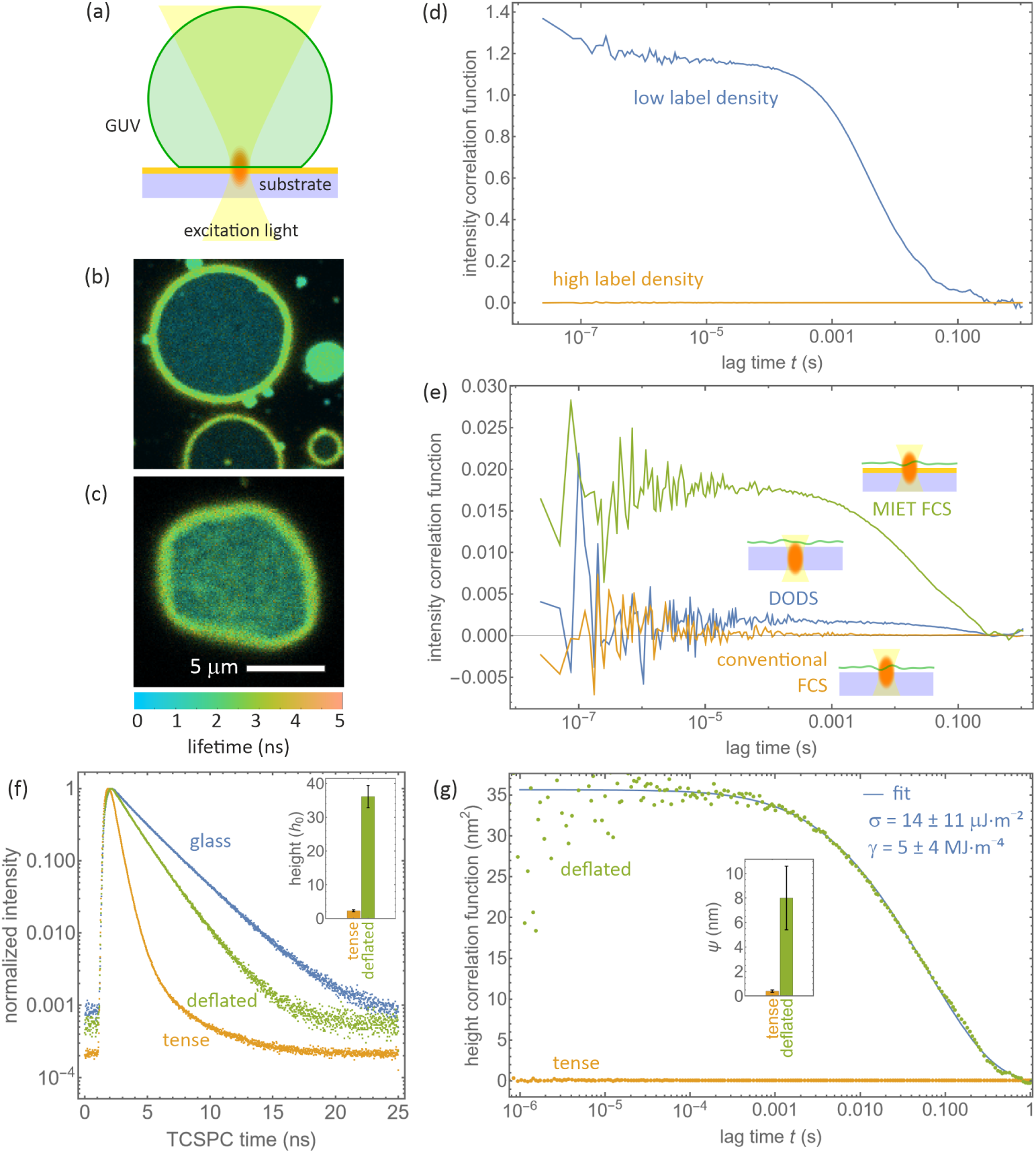
(a) Schematic of a MIET-FCS experiment on a GUV attached to the MIET substrate. (b) Fluorescence lifetime image of a tense GUV showing a ring of increased intensity where the GUV membrane first touches the surface. Fluorescence lifetime is shown by color, intensity by brightness. (c) Same as (b), but for a deflated GUV. (d) Intensity correlation curves measured on the proximal membrane of GUVs immobilized on a pure glass substrate. Shown are two curves for membranes with low and high label density. (e) Comparison of correlation curves measured on the proximal membrane of surface-immobilized GUVs obtained on pure glass substrate with the excitation focus on the glass surface (conventional FCS) or ca. 1 µm below (DODS), and obtained on a MIET substrate. (f) Fluorescence lifetime decay curves for tense GUVs on glass and MIET substrate, and for a deflated GUV on MIET substrate. (g) Measured MIET-FCS correlation curves (dotted lines) for tense and deflated GUVs. Inset presents corresponding height fluctuation amplitudes (*ψ*). Fit values from a fit (solid line) of eq. 4 to the deflated GUV curve are also shown.

Next, we performed proof-of-principle experiments to show that MIET-FCS is indeed capable of significantly amplifying the amplitude of minute membrane fluctuations that cannot be observed by FCS on glass substrates with GUVs, and are barely visible when positioning the focus slight below the glass-water interface as in a DODS experiment. In particular, we immersed GUVs (230 mOs/mL) into a hypertonic solution with an osmolarity of 400 mOs/mL, which deflates the GUVs and leads to strong membrane fluctuations. The deflation is clearly visible in the non-circular geometry of the contact area of a GUV settled on a BSA-covered glass surface, shown in Figure 2b. In Figure 2e we show the comparison of three FCS measurements with the following arrangements: GUV on glass with the excitation focus located at the substrate surface; GUV on glass with the excitation focus positioned much below the glass surface (as used in DODS experiments); GUV on a MIET substrate with the excitation focus on the substrate surface. As anticipated, we did not observe any lag-time-dependent correlation decay for the GUV on glass when the membrane was in focus, and we observed a very small decay amplitude (*<* 10*^−^*^3^) for the GUV on glass but with defocused excitation. In contrast, the FCS measurement on a MIET substrate yields an correlation function with considerable amplitude (∼ 0.02), that is an order of magnitude larger. Moreover, from the DODS FCS measurement, it is difficult to derive a precise number for the root mean square height fluctuation since this value is extremely sensitive on the exact vertical position of the laser focus. In contrast, using the mean lifetime values of the GUV measured on the MIET substrate, we can calculate a mean height value, which in this example is about *h*_0_ ∼ 37.0 nm. This allows us to use eq. (3) to determine the value of the root mean square height fluctuation *ψ* of 6.0 nm from the amplitude of the height correlation curve.

Next, we used MIET FCS to compare the membrane fluctuations of tense GUVs (immersed in an isotonic buffer with 230 mOs/mL) and deflated GUVs (immersed into a buffer with of 400 mOs/mL). Figure 2f shows three TCSPC measurements of the fluorescence decays for a tense GUV on pure glass, on a MIET substrate, and for a deflated GUV on a MIET substrate. One can clearly see the MIET induced reduction of the fluorescence lifetime as compared to the pure-glass measurement for both, tense as well as deflated GUVs. Moreover, a tense GUV on a MIET substrate shows a much shorter fluorescence lifetime as compared to a deflated GUV on the same substrate (0.49 ns for tense GUV and 1.43 ns for deflated GUV). We converted these fluorescence lifetimes into height values, which yields *h*_0_ = 2.3 ± 0.3 nm (*N* = 23) for tense GUVs, and *h*_0_ = 36.1 ± 3.3 nm (*N* = 36) for deflated GUVs, which signifies that the average height of the basal membrane of the tense GUVs is much smaller than that of the deflated GUVs (see inset of Figure 2f). Figure 2g shows a comparison of the height correlation curves *g_h_* for the tense and deflated GUVs on the MIET substrate. As can be seen, the height fluctuations for the tense GUV are nearly negligible (0.39±0.11 nm, *N* = 23), while the deflated GUV shows by orders of magnitude larger fluctuations with a root mean square height fluctuation of *ψ* = 8.0±2.6 nm (*N* = 36), see also inset of Figure 2g. This is in excellent agreement with published values, such as obtained with FLIC microscopy (*h*_0_ = 30-60 nm and *ψ* = 10 nm) [16–18], or RICM (*h*_0_ = 31-39 nm and *ψ* = 4-15 nm) [42]. The relaxation time *τ^∗^*, defined as the lag time at which the height correlation curve has decreased to half its maximum value, was determined to be 50 ± 30 ms using MIET FCS for the deflated membrane, which is consistent with previous findings in the literature [14].

We fitted the measured height correlation function *g_h_*(*t*) with a theoretical model described in Refs. [42–45]. The fluctuations are determined by the membrane bending rigidity *κ*, the membrane tension *σ*, the effective viscosity *η*, and an effective interaction potential *γ* describing the interaction between the fluctuating membrane and the substrate surface. Then, the height correlation function is described by the following equation:

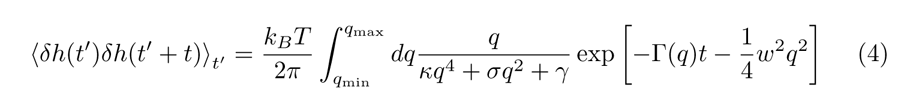

where *w* denotes the diameter of the confocal detection area. The function Γ(*q*) is defined by

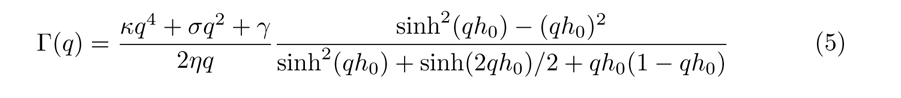

The integration boundaries of the integral in eq. (4) are *q*_min_ = (*γ/κ*)^1*/*4^ and *q*_max_ = 1*/h*_0_ [45].

For SOPC membranes, the value of the bending modulus *κ* = 20 *k_B_T* is well known and was kept constant throughout our analysis [14, 46]. The effective viscosity *η* = 1.2 mPas was calculated as the arithmetic mean of the viscosity of the buffer solutions inside and outside the GUV. The diameter of the confocal detection area in our measurement was determined as *w* = 280 nm. We measured and fitted *N* = 36 height fluctuation correlation curves, a typical curve and fit is shown in Figure 2g. From these fits, we found a surface tension value of *σ* = 14 ± 11 µJm^-2^ and an interaction potential strength value of *γ* = 5 ± 4 MJm^-4^, in excellent agreement with published results for similar GUV measurements [42–44].

#### 3.1.2 Membrane fluctuations of red blood cells

We applied MIET FCS to a well-studied biological system by measuring membrane fluctuations in red blood cells (RBCs). It is well-known that RBC membranes exhibit significant bending fluctuations, which have been extensively studied in the past using various techniques. These include studies conducted by Park et al. [3] and Betz et al. [4], as well as Monzel et al. [14, 47] and Gov et al. [47].

When RBCs settle on a planar surface, the contact zone between their membrane and the surface forms a circular rim-like structure, owing to the concave shape of these cells. This can be seen in Figures 3a,b. A high label density is used here again to ensure that the lateral diffusion did not affect our measurement (see Supplementary Figure 3).

**Fig. 3.**
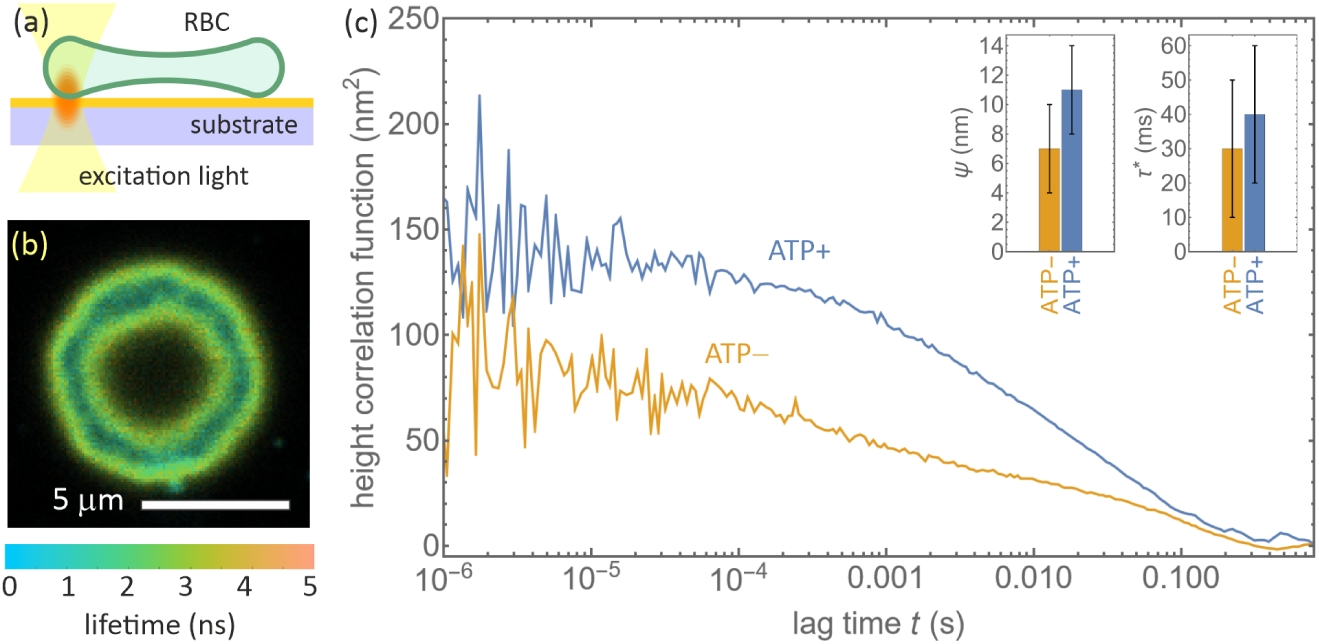
(a) Schematic of the MIET-FCS experiment of membrane fluctuations in a living RBC. An RBC has settled on the BSA covered surface of a MIET substrate. (b) Fluorescence lifetime image of a RBC on a MIET substrate. Lifetime is shown by color, fluorescence intensity by brightness. The cell’s rim touching the surface is visible as a ring of enhanced fluorescence intensity and reduced lifetime. (c) Height fluctuation correlation curves measured on the membrane of an ATP-saturated and an ATP-depleted RBC. Insets show histograms of fluctuation amplitudes (*ψ*) and relaxation times (*τ ^∗^*). Error bars denote standard deviations (*N* = 18 for ATPand *N* = 14 for ATP+ measurements).

We conducted MIET-FCS measurements on the proximal membrane of the circular rim formed by RBCs settled on a BSA-coated MIET substrate. This enabled us to measure the membrane fluctuations in the presence and absence of ATP, as shown in Figure 3c. The average height values of the membrane did not appear to be affected by the RBC’s biological activity, with values of 36 ± 6 nm in the absence of ATP and 37 ± 5 nm in the presence of ATP.

However, we observed that membrane fluctuation amplitudes were larger in the presence of ATP, with an increase of 1.6 times as compared to ATP-depleted RBCs. Specifically, the amplitude values were 7 ± 3 nm and 11 ± 3 nm in the absence and presence of ATP, respectively. Additionally, the relaxation time was measured to be *τ^∗^* = 30 ± 20 ms for ATP-depleted cells, which was slightly smaller than the value of 40 ± 20 ms in the presence of ATP. This suggests a faster dissipation of thermal-driven fluctuations for active cells, in good agreement with previously published results [14].

### 3.2 GIET-FCS

The previous two subsections employed MIET, a technique based on the metal-induced quenching of fluorescence, to measure membrane fluctuations in GUVs and red blood cells. In the following two subsections, we will use GIET, which relies on the energy transfer from excited fluorescent dyes to excitons in a single sheet of graphene.

The dynamic range of GIET is approximately 8 times smaller than that of MIET, at around 20 nm compared to 160 nm. However, this limitation allows GIET to achieve greater sensitivity to vertical motions of an emitter, making it ideally suited for measuring extremely small fluctuations down to the ^°^Angström length scale.

#### 3.2.1 Fluctuations of a pore spanning membrane

SLBs are a gold standard for *in vitro* studies of lipid membranes. However, the direct contact between the membrane and the solid support can significantly alter the dynamics and properties of SLBs. Therefore, it is important to be able to study free-standing membranes that are not disturbed by solid supports. One way for achieving this is using pore-spanning membranes (PSMs), where a membrane is formed over small pores in a solid substrate, as shown in Figure 4a.

**Fig. 4.**
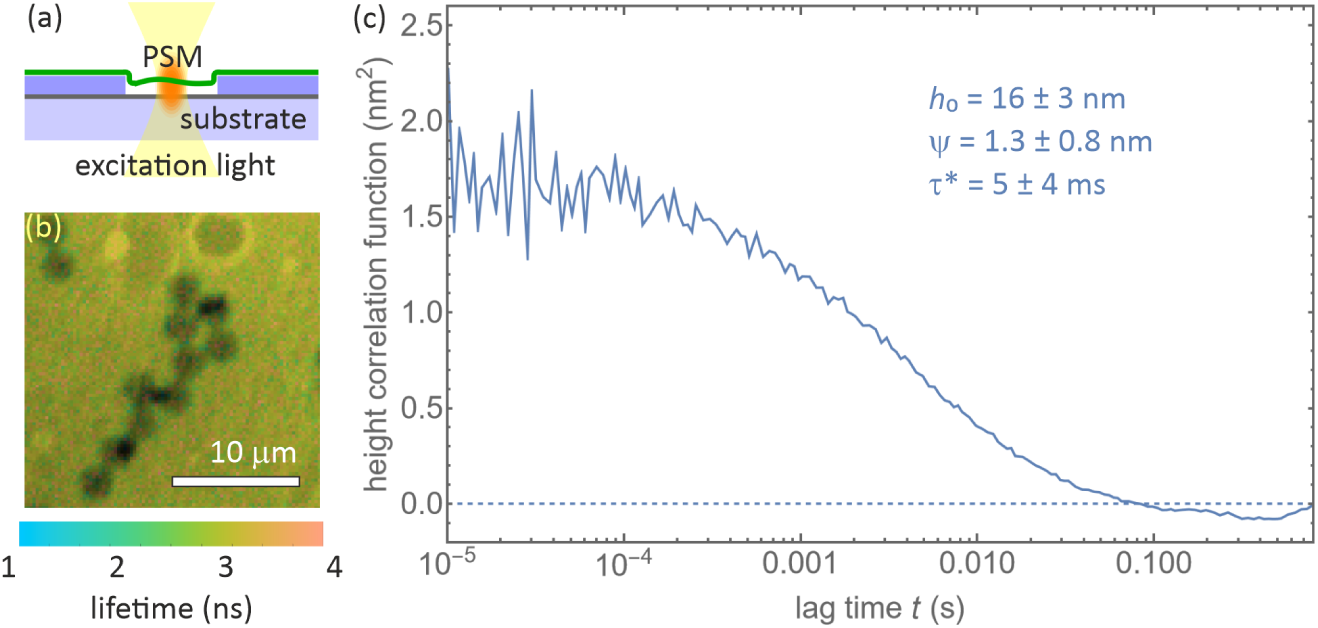
(a) Schematic of a GIET-FCS experiment on a PSM. The GIET substrate consists of a single sheet of graphene sandwiched between a glass cover slide and a 32 nm thin SiO_2_ spacer layer. Into that SiO_2_ layer, pores were etched with a depth of 30 nm and a diameter of 3 µm. (b) Fluorescence lifetime image of a PSM sample. Lifetime is shown by color, fluorescence intensity by brightness. Several pores are visible as dark disks on a brighter background. Two types of pores with different intensity/lifetime can be distinguished, corresponding to free-standing membranes and membranes touching the pore bottom (reduced intensity/lifetime). (c) Height fluctuation correlation curves for a PSM on a GIET substrate.

Compared to GUVs, PSMs typically have a much larger surface tension, which reduces their transversal fluctuations by orders of magnitude [48, 49]. Currently, no technique has been reported that can measure such small fluctuations in PSMs.

We utilized pore-spanning membranes (PSMs) to prepare free-standing membranes away from a solid support, which can be important when studying membrane properties that may be affected by contact with a solid support. To create PSMs, we deposited negatively charged giant unilamellar vesicles (GUVs) onto a positively charged substrate with pores (see Method and Supplenentary Figure 6) [50]. This substrate consisted of a single sheet of graphene sandwiched between a glass cover slide and a thin SiO_2_ spacer layer with a thickness of 32 nm, into which pores of 3µm diameter and 30 nm depth were etched, as shown in Figure 4b. After deposition, the GUVs ruptured to form PSMs.

We confirmed the free-standing nature of the PSMs using GIET and GIET-FCS by measuring their average height and fluctuation amplitudes (see Supplementary Figure 5 and Supplementary Note 2). The fluorescence lifetime image in Figure 4b clearly distinguishes pores showing a small and long lifetime, representing membranes touching the pore bottom and free-standing membranes, respectively. From lifetime measurements, we found an average height of 16.3 ± 2.6nm (*N* = 20) for the free-standing membranes, indicating that the membrane bends into the pore due to membrane prestress [51]. We performed GIET-FCS measurements at the center of the PSMs, as shown by the solid line in Figure 4c. As a control, we performed similar measurements on PSMs over pores without a graphene sheet (see Supplementary Figure 5a). As expected, the control measurement did not show any intensity correlation amplitude or decay, in stark contrast to the GIET-FCS measurements (see Supplementary Figure 6). In the latter measurements, we observe a height correlation amplitude of 1.3 ± 0.8 nm (*N* = 20) and a correlation relaxation time of *τ^∗^* = 4.5 ± 3.7 ms. However, the height fluctuation correlation curve of PSMs cannot be well fitted with the theoretical model used for GUV fluctuations in section 3.1.1, due to the more complex hydrodynamic interactions of the fluctuating membranes with the liquid enclosed in the membrane-covered pore.

We did not detect any significant fluctuations using GIET-FCS when the pore depth was reduced to 20 nm (see Supplementary Figure 6). In this case, the PSMs are too close to the pore bottom, with an average height of 8.2 ± 1.3 nm, as determined from lifetime measurements. This proximity to the substrate dramatically reduces the amplitude of membrane fluctuations.

#### 3.2.2 Fluctuations of mitochondrial membranes

Mitochondria are dynamic organelles that play a vital role in cellular energy production. They are characterized by a double-membrane structure [52]. The smooth outer mitochondrial membrane (OMM) surrounds the organelle and contains numerous protein-based pores, while the inner mitochondrial membrane (IMM) is deeply convoluted and contains the machinery responsible for ATP synthesis [52]. It has been shown that isolated mitochondria can maintain their biological activity, such as their ability to synthesize ATP, in the presence of suitable precursors in the surrounding buffer [53].

To investigate the effect of ATP-synthesis precursor molecules on mitochondrial membrane dynamics, we performed GIET-FCS experiments on isolated mitochondria in two different states: an ”active” state with a buffer containing ADP molecules that promote ATP synthesis, and a ”resting” state with a buffer lacking these molecules (see subsection 5.8 in Materials and Methods). To label specific regions of the mitochondria, we attached fluorescent probes to either the outer mitochondrial membrane (OMM) or the inner mitochondrial membrane (IMM), as shown in Figures 5a,b. To ensure that lateral diffusion did not affect our measurements, we used a label density that was sufficiently high (see Supplementary Figure 7 and Supplementary Note 3).

**Fig. 5.**
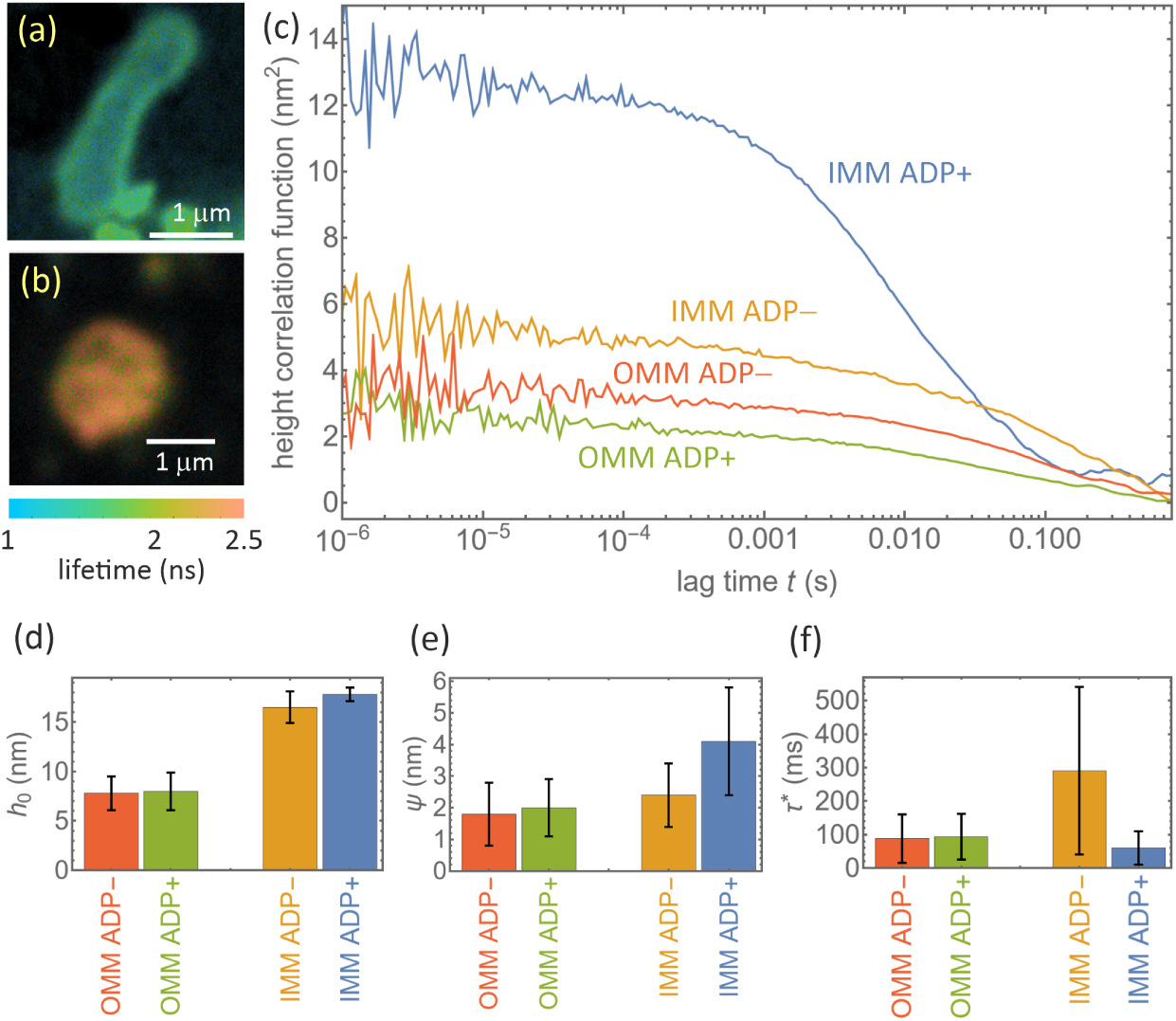
(a) FLIM image of the OMM above a graphene substrate. (b) Same as (a) for the IMM. (c) Height correlation curves for the IMM and OMM under ADP+ and ADPconditions. (d) Average height values (*h*_0_). (e) Height correlation amplitude, *ψ*. (f) Correlation relaxation times, *τ ^∗^*. In (d-f), error bars denote standard deviations.

We used a GIET substrate consisting of a single layer of graphene on a glass cover slide, with a 10 nm thick SiO_2_ spacer layer for OMM measurements and a 2 nm thick SiO_2_ spacer layer for IMM measurements.

We conducted comparative GIET-FCS experiments on isolated mitochondria using a buffer containing ATP-synthesis precursor molecules (ADP+ active state) and a buffer without these molecules (ADPresting state). For OMM measurements, we used a GIET substrate composed of a single sheet of graphene deposited on a glass cover slide and covered with a 10 nm thick SiO_2_ spacer layer, while for IMM measurements, we used a 2 nm thick SiO_2_ spacer layer. We measured *N* = 22 mitochondria under ADPand *N* = 21 mitochondria under ADP+ conditions for OMM, and *N* = 21 under ADPand *N* = 26 under ADP+ conditions for IMM. Exemplary GIET-FCS curves are shown in Figure 5c, and results for the average distance, height fluctuation amplitudes, and correlation relaxation times are presented in Figures 5d-f.

Our results showed that the average membrane height *h*_0_ for OMM remained almost constant for mitochondria in the resting (7.8±1.7 nm) and active (8.0±1.9 nm) states. However, this value decreased by about 1.3 nm for IMM (16.5 ± 1.6 nm for ADP-resting and 17.8 ± 0.7nm for ADP+ active states), consistent with previous reports using static GIET measurements [37]. Using GIET-FCS, we found a similar trend for the height fluctuation amplitudes: OMMs exhibited comparable fluctuation amplitudes *ψ* for mitochondria in their resting and active states (1.8±1.0 nm for ADP-resting and 2.0 ± 0.9 nm for ADP+ active states), while IMMs displayed enhanced fluctuation amplitudes in active state mitochondria (2.4 ± 1.0 nm for ADP-resting and 4.1 ± 1.7 nm for ADP+ active states).

Finally, the correlation relaxation time (*τ^∗^*) also exhibited a difference between the two membrane types. For IMMs, the values decreased from 0.29 ± 0.25 in the resting state to 0.06±0.05 in the active state, while for OMMs, almost no change was observed (0.088 ± 0.072 for the ADP-resting and 0.093 ± 0.068 for the ADP+ active states). These results suggest that the active membrane fluctuations of IMMs are enhanced in the mitochondria’s active state, likely due to structural reorganization of the IMMs’ cristae and mitochondrial volume regulation, as reported previously [37].

## 4 Discussion

We have developed a novel method, MIET/GIET-FCS, which enables the study of lipid membrane fluctuations. This technique combines the nanometer spatial resolution of MIET/GIET with the microsecond temporal resolution of FCS. What sets our approach apart is its simplicity, as it can be implemented using a conventional fluorescence lifetime imaging confocal microscope. Additionally, our method eliminates the need for experimental calibration by providing exact MIET/GIET calibration curves based on sample geometry.

Through the application of MIET/GIET-FCS to various systems, including GUVs, living red blood cells, pore-spanning membranes, and the inner and outer membranes of living mitochondria, we have demonstrated its versatility for a wide range of membrane biophysical studies. By expanding MIET/GIET-FCS to a large-field of view using a combination of a widefield microscope and a fluorescence lifetime camera, we can achieve high-statistical power measurements of cell membrane fluctuations. This technique holds great promise for assessing cellular functioning in response to drugs and therapies, providing insights into health, mechanical properties, and treatment responses for improved healthcare outcomes. Our method has vast potential for applications in the study of biological membranes both *in vitro* and *in vivo*.

## 5 Materials and Methods

### 5.1 MIET and GIET measurements

All measurements, including lifetime measurements, were conducted using a self-constructed confocal setup. We used a pulsed diode laser (LDH-D-C 640, PicoQuant) with a pulse width of 50 ps FWHM and a repetition rate of 40 MHz, operating at an excitation wavelength of 640nm. The excitation light was filtered using a clean-up filter (LD01-640/8 Semrock) and collimated through an infinity-corrected 4x objective (UPISApo 4X, Olympus). The laser beam was then directed to a high numerical aperture objective (UApoN 100X, oil, 1.49 N.A., Olympus) via a dichroic mirror (Di01-R405/488/561/635, Semrock). The emission light was collected and focused into a 100 µm diameter pinhole and refocused onto two avalanche photodiodes (*τ* - SPAD, PicoQuant). We utilized a long-pass filter (BLP01-647R-25, Semrock) and two band-pass filters (Brightline HC692/40, Semrock) before the pinhole and the detector, respectively. The photo signals obtained from the detector were analyzed using a multi-channel picosecond event timer (Hydraharp 400, PicoQuant) with 16 ps time resolution.

In a typical measurement, we first scanned the sample at the coverslip surface to determine the contour of the GUV, cell, or mitochondrion and selected the appropriate point position for data acquisition. The laser power was adjusted to reach a maximum count rate of 50-250 kcps during data collection. For GUV, mitochondrion, and cell measurements, data were recorded for at least 3 minutes, while for the pore-spanning membrane, at least 5 minutes were used to build the intensity correlation curve. For planar membrane measurements with dense labeling (SLB and tense GUV), at least 20 minutes were used to smooth the correlation curve.

To analyze the data, we used a custom-written MATLAB-based software package that includes the calculation of MIET lifetime-versus-distance curves and the conversion of lifetime to distance. This software is equipped with a graphical user interface and has been published [54]. It is available free of charge at https://projects.gwdg.de/ projects/miet.

### 5.2 Substrate preparation

The protocol for preparing the gold-modified substrate has been previously described in our publications [26, 28]. Briefly, we used a layer-by-layer electron-beam evaporation process to deposit a 2 nm titanium layer, followed by a 10 nm gold layer, another 1 nm titanium layer, and finally a 10 nm SiO_2_ layer on the surface of a glass coverslip. This gold-covered substrate is referred to as the MIET substrate. Prior to conducting MIET experiments, the MIET substrate was subjected to plasma cleaning for 60 seconds at the highest plasma intensity (Harrick Plasma). Subsequently, it was incubated in 5 mg/mL^-1^ bovine serum albumin (BSA, Sigma-Aldrich) solution for 15 minutes to prevent any non-specific interactions during experiments involving GUVs and RBCs. For GIET, we used a transfer method based on the manufacturer’s instructions (Easytransfer, Graphena. Inc.) to prepare a graphene-modified substrate. Silica with the desired thickness was then evaporated onto the surface of the graphene-modified substrate to create the GIET substrate. To prepare the pore-containing graphene substrate, we employed the nanosphere lithography method (Supplementary Figure 4) [55]. First, we evaporated a 2 nm SiO_2_ layer onto the graphene-modified substrate to protect the graphene film. Then, we deposited polystyrene (PS) beads with a diameter of 3 µm (Bangs Laboratories, Inc.) in water onto the SiO_2_/graphene substrate. This substrate was then dried at room temperature. Next, silica with the desired thickness (30 nm) was evaporated on top of the PS/SiO_2_/graphene substrate. Finally, the PS beads were removed in acetone under ultrasonication for 5 minutes, resulting in a substrate with pores of the desired height and a diameter of 3 µm on the graphene surface.

For the pore substrate without graphene for pore spanning membrane (PSM) measurements above the glass, we deposited the PS beads onto the coverslip and coated all layer-by-layer with 2 nm Ti, 10 nm Au, 1 nm Ti and 20 nm SiO_2_. The gold film was used to quench the fluorescence of the supported membrane, allowing us to distinguish the PSMs from the GUV patch. Before using the substrates for membrane spanning, we modified the substrates to have a positively charged surface [48]. We first treated the substrates with plasma cleaning for 30 seconds at low intensity and then incubated them in a solution of 1% (v/v) (3-trimethoxysilylpropyl)-diethylenetriamine (DETA) in water for 15 minutes. The positively charged substrates were then washed with methanol and water, followed by heating at 110*^◦^*C for 15 min.

To prepare the graphene substrate for mitochondria measurements, we used a 2 nm or 10 nm thick SiO_2_-spacer and treated it with 0.1% (wt/v) poly-L-lysine (PLL) solution for 5 minutes. The substrate was then rinsed with water and allowed to dry. The resulting positively charged substrates were stored in a clean, dust-free environment to prevent contamination.

### 5.3 Lipids

1-stearoyl-2-oleoyl-sn-glycero-3-phosphocholine (DOPC), 1,2-dioleoyl-sn-glycero-3-phosphoethanolamine-N-(methoxy (polyethyleneglycol)-2000) (DOPE-PEG2000), (1,2-dioleoyl-sn-glycero-3-phosphoethanolamine-N-(cap biotinyl) (DOPE-cap-biotin), 1,2-dioleoyl-sn-glycero-3-phosphoethanolamine (DOPE), 1,2-dioleoyl-3-trime thylam-moniumpropane,chloride salt (DOTAP), 1,2-diphytanoyl-sn-glycero-3-phosphocholine (DPhPC) and 1,2-dioleoyl-sn-glycero-3-phospho-(1’-rac-glycerol) (DOPG) were purchased from Avanti Polaer Lipids and diluted to 10 mg/mL in chloroform as stock solution and stored at -20℃. The Atto655-dyes labeled 1,2-dipalmitoyl-sn-glycero-3-phosphoethanolamine (DPPE-Atto655) were purchased from ATTO-TEC GmbH and dissolved in chloroform at a concentration of 0.01 mg/mL and 1 mg/mL for stock solution.

### 5.4 Vesicles preparation

Giant unilamellar vesicles (GUVs) were produced using the electro-swollen method [28]. For MIET measurements, a lipid mixture containing SOPC with 2 mol% DOPE-PEG2000, 5 mol% DOPE-cap-biotin and 0.001-1 mol% DPPE-Atto655 was used, while for PSM measurements, a mixture containing DPhPC with 10 mol% DOPG and 0.001-1 mol% DPPE-Atto655 was used. The mixture was deposited onto an electrode and allowed to evaporate under vacuum for 3 hours at 30*^◦^*C. The chamber was then filled with a sucrose solution (230 mM for MIET-GUV measurements and 300 mM for PSM measurements). An alternating electric current at 15 Hz and a peak-to-peak voltage of 1.6 followed was applied for 3 hours to the chamber used for electro-formation, followed by a treatment with 8 Hz for 30 minutes. The final GUV suspension was stored at -4*^◦^*C and used within 3 days.

For the MIET-GUV experiments, the GUV suspension was diluted 50-folded in PBS buffer (10 mM Na_2_HPO_4_, 2 mM KH_2_PO_4_, and 3 mM KCl, pH 7.4) with varying osmolarity by adding NaCl. Deflated GUVs were observed in PBS buffer with 400 mOsml^-1^, while tense GUVs were observed in PBS buffer with 230 mOs/mL. For PSM experiments, the stock GUV suspension was diluted 50-fold in Tris-HCl buffer (20 mM Tris-HCl, 100 mM NaCl, 10 nM CaCl_2_, pH 7.4). For all the GUV experiments, the diluted GUV solution with observation buffer was added to a Culture-Insert 4 Well chamber (ibidi GmbH) on the modified substrate. The chamber was then covered with a glass slide and kept at room temperature for 15 minutes to allow for equilibration before measurement.

### 5.5 Preparation of RBCs

Human RBCs were prepared from a healthy donor (author Tao Chen) by pricking a finger and diluting the blood 100-fold with PBS buffer (osmolarity of 300 mOs/mL). The resulting suspension was collected by centrifugation at 200 g for 1 minute, and the pellet was fluorescently stained using a liposome method (see next subsection). The stained pellet was then washed twice with PBS buffer and dissolved in the observation buffer. For the ATP-saturated RBC experiments, the observation buffer contained 0.1 mg/mL BSA and 10 mM D-Glucose, while for the ATP-depleted RBCs, the buffer contained only 0.1 mg/mL BSA. The RBCs were ATP-depleted by incubating them in the observation buffer overnight at 37*^◦^*C. After staining, the cells were used within one hour.

### 5.6 Fluorescent staining of RBCs

We employed a fusogenic liposome method [14, 56] to label the RBCs with a fluorescent marker. Initially, a lipid mixture comprising DOPE, DOTAP, and DPPE-Atto655 in a 1:1:0.2 weight ratio was dried under vacuum for 3 hours at 30*^◦^*C and re-suspended in 20 mM Tris-HCl (pH 7.4) buffer at a concentration of 2.2 mg/mL. The suspension was vortexed for 2 minutes and sonicated for 10 minutes to generate small multilamellar liposomes. To stain the cells, RBCs were incubated in the diluted liposome solution (1:100) for 15 minutes at 37*^◦^*C. The resultant fluorescently labeled RBCs were then washed twice with PBS buffer (300 mOsml^-1^) by centrifugation (200 g, 1 min) and re-suspended in the observation buffer.

### 5.7 Isolation of mitochondria

The isolation of mitochondria from cultured MH-S cells was performed using the Thermo Scientific™ mitochondrial isolation kit. The isolated mitochondria were snap-frozen in liquid nitrogen and stored at -80*^◦^*C in 300 mM trehalose buffer (300 mM trehalose, 10 mM HEPES-OH, 10 mM KCl, 1 mM EGTA, and 0.1% BSA, pH 7.7). This buffer has been reported to preserve the biological functions of mitochondria and maintain the integrity of their outer membrane [57].

### 5.8 Mitochondria membranes measurement

In a typical measurement of mitochondria, 5 µL of frozen mitochondria were thawed by holding the tube between fingers and mixed with 25 µL of trehalose buffer. The resulting solution was added on top of the PLL-modified graphene substrate with a 2 nm SiO_2_ spacer for IMM or a 10 nm SiO_2_ spacer for OMM measurements. The plates were incubated at 4*^◦^*C for 30 minutes, followed by the addition of 70 µL of respiration buffer [53] (130 mM KCl, 5 mM K_2_HPO_4_, 20 mM MOPS, 1 mM EGTA, 0.1% (w/v) BSA, pH 7.1) containing MitoTracker™ Deep Red (500 nM) or CellMask™ Deep Red (10 µg/mL). After 15 minutes of incubation at room temperature, the buffer was carefully changed to a new respiration buffer without dye. For ADP stimulated respiration measurements, we added 1 mM malic acid, 5 mM pyruvate, and 100 µM ADP to the respiration buffer. The mitochondria were used for up to one hour after fluorescent staining.

## Supplementary information

If your article has accompanying supplementary file/s please state so here.

## Acknowledgments

We thank Anna Chizhik, Ingo Gregor and Alexey Chizhik their help and support.

## Declarations

- Funding: T.C. and J.E. acknowledge financial support by the European Research Council (ERC) for financial support via project “smMIET” (grant agreement no. 884488) under the European Union’s Horizon 2020 research and innovation program. J.E. acknowledges financial support by the DFG through Germany’s Excellence Strategy EXC 2067/1-390729940.
- Conflict of interest: The authors declare no conflict of interest.
- Availability of data and materials: The data for all curves of Figures 1-5 are deposited on GitHub at https://gitlab.gwdg.de/igregor/ miet-membrane-fluctuation.git. A detailed description is given in the file Inventory.pdf.
- Code availability: All code used for generating Figures 1-5 are deposited on GitHub at https://gitlab.gwdg.de/igregor/miet-membrane-fluctuation.git. In particular, the depository contains *Matlab* (MathWorks^®^ Inc.) code used for calculating the model curves of Figure 1c,d and for fitting the deflated GUV height correlation curve in Figure 2g, and a *Mathematica* (Wolfram Research Inc.) notebook that generates the graphs of all Figures.
- Contributions of authors: T.C. prepared all samples, performed all the measurements. T.C. and N.K. and analyzed the data. J.E. made all figures in the main text. All authors helped with preparing the manuscript, which was written by J.E.

---

**Supplementary Figure 1:**
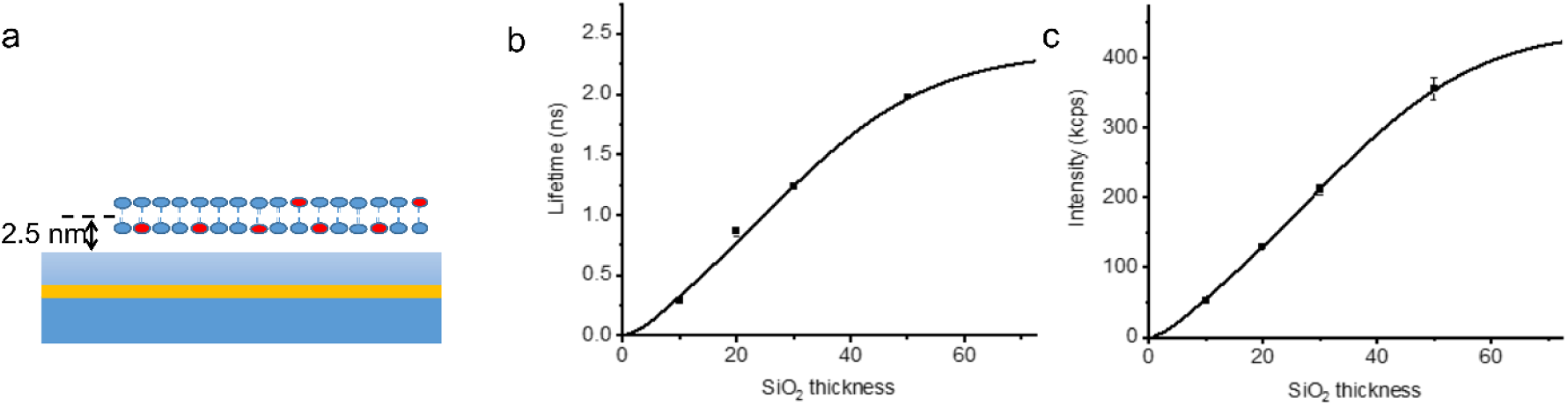
Experimental verification of theoretical calibration curves. (a) Schematic of a MIET calibration experiment used to validate the theoretical calculations: a glass cover slide is covered with a 20 nm thick gold layer that is covered with SiO2 spacer layer of varying thickness. On top of that, a supported lipid bilayer (SLB) with fluorescently labeled lipids is prepared. In panels (b) and (c), the solid curves represent calculated MIET lifetime and brightness curves, respectively, for an electric dipole with an emission wavelength of 680 nm, a quantum yield of 0.36, a free space lifetime of 2.6 ns. The refractive index of the dipole-embedding medium (aqueous buffer solution in experiments) was set to 1.33, and that of the SiO2 spacer to 1.46. The experimental data in panels (b) and (c) were measured with SLBs comprising DOPC and 0.001% DPPE-Atto655 on four substrates with different silica spacer thicknesses (10 nm, 20 nm, 30 nm, and 50 nm). The error bars represent the standard deviations of the measurements.

**Supplementary Figure 2:**
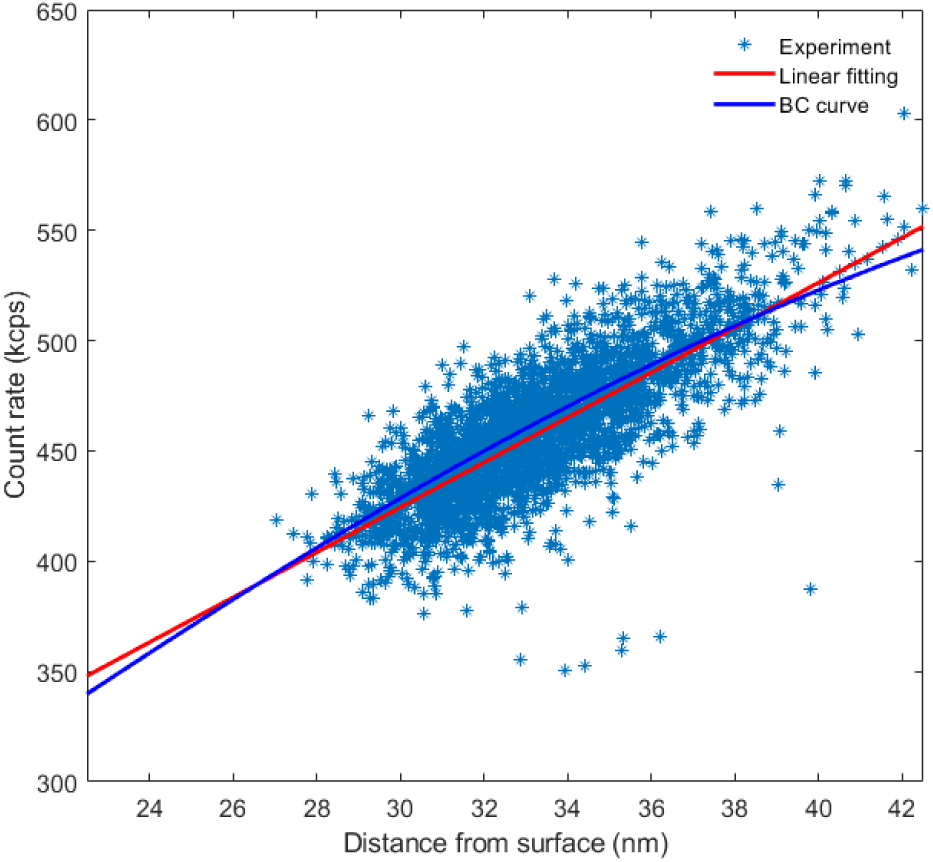
Check of linearity for the relation between height and fluorescence intensity. Scatter plot showing the correlation between measured intensities and respective height values as calculated from fluorescence lifetimes, as obtained for a deflated GUV. The experiment lasted for 300 s and was divided into 3000 bins of 100 ms time width. The red line represents a linear fit of the scatter plot data, and it has a slope of 10.2 kcps/nm. The blue line represents the theoretical brightness calibration curve, having a slope at a distance of 33 nm of 9.8 kcps/nm.

**Supplementary Figure 3:**
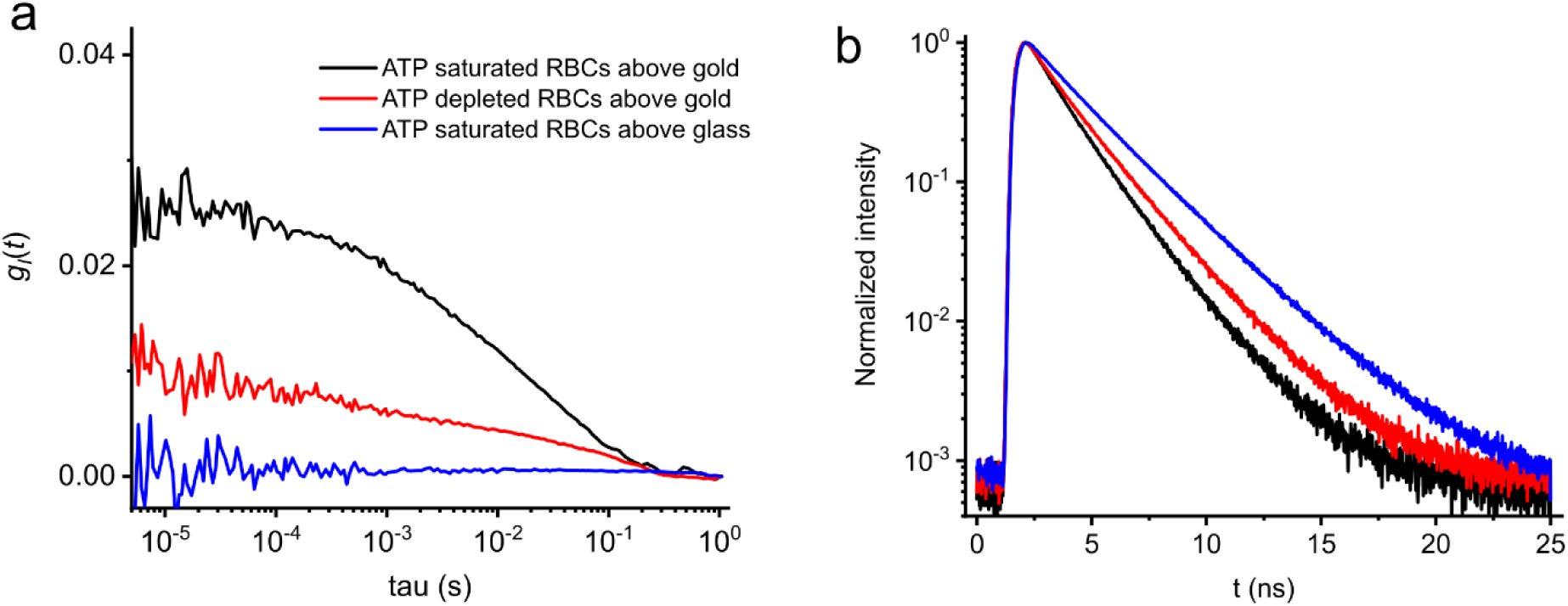
Intensity correlations and lifetime decay curves for RBCs. (a) Intensity autocorrelation function, *gI*(*t*), measured at the rim of RBCs on different substrates and for ATP-saturated and ATP-depleted buffers. (b) Corresponding normalized fluorescence lifetime curves. The intensity correlation measurement at the glass surface showed almost no amplitude, indicating that the dye concentration used was sufficiently large to suppress any diffusion-related contribution to the correlation curve.

**Supplementary Figure 4:**
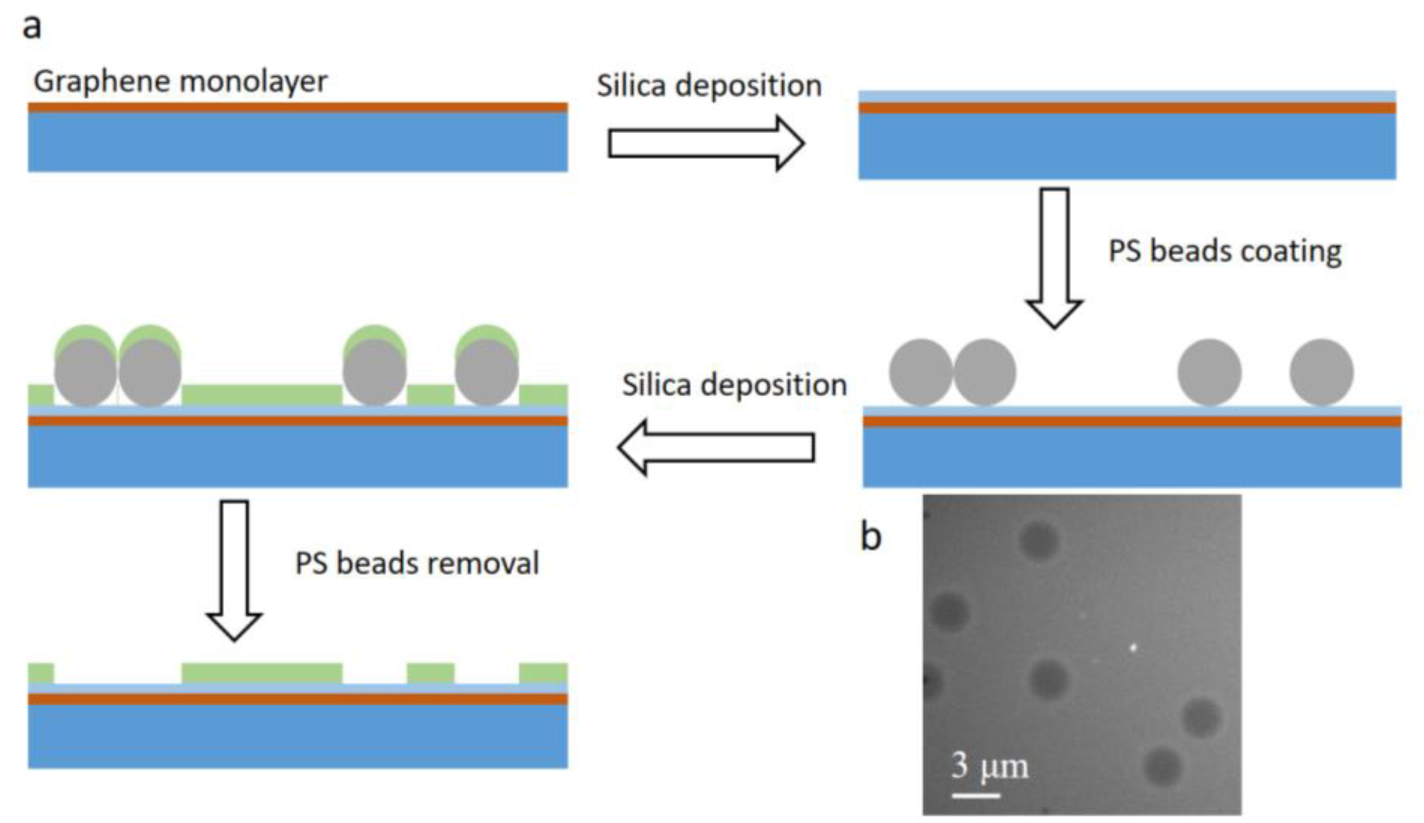
Fabrication of graphene-pore substrate. (a) Schematic of GIET substrate fabrication for PSM measurements. (b) SEM images of the pore-covered GIET substrate.

**Supplementary Figure 5:**
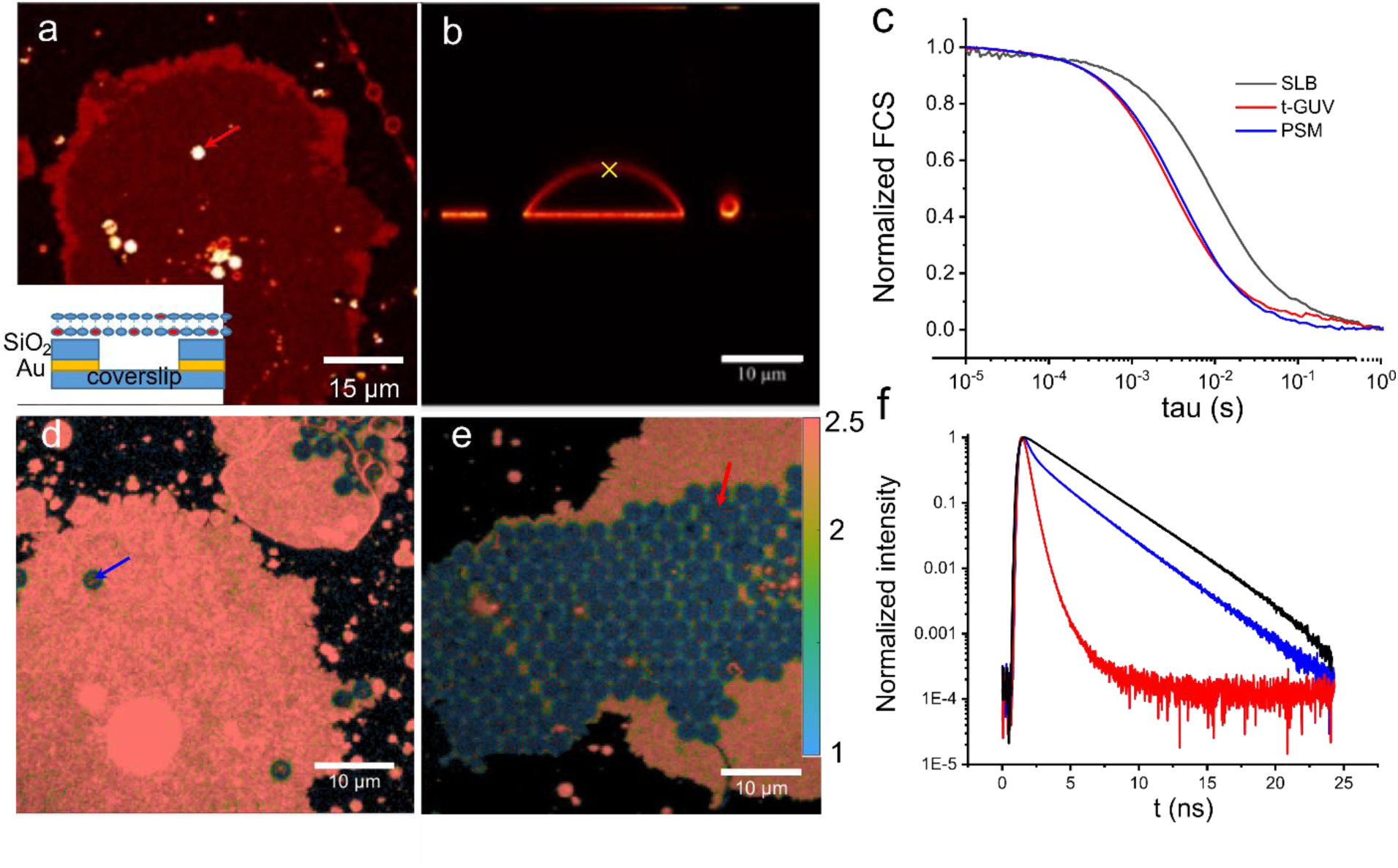
Checking the generation of free-standing membranes. (a) Confocal fluorescence image showing a GUV patch on a glass-gold substrate, with an arrow indicating the position of the PSM above the glass used for FCS measurements. The inset shows a schematic representation of the structure of the substrate. (b) Confocal fluorescence image showing a vertical section of a GUV on a glass/BSA substrate, with a cross indicating the position at the top of the membrane used for FCS measurements (t-GUV). (c) FCS curves of DPPE-Atto655 for SLB, t-GUV, and PSM above glass. (d) Confocal fluorescence lifetime image showing a GUV patch on a GIET substrate, with an arrow indicating the position used for the PSM lifetime measurement. (e) Confocal fluorescence lifetime image of a GUV patch on a multi-pore graphene substrate, with an arrow indicating the position used for the lifetime measurement. (f) Fluorescence lifetime curves for a SLB on glass, for a membrane on the bottom of a pore (t-GUV), see red arrow in (e), and a PSM over a pore with 30-nm depth above a graphene substrate, see blue arrow in (d).

**Supplementary Figure 6:**
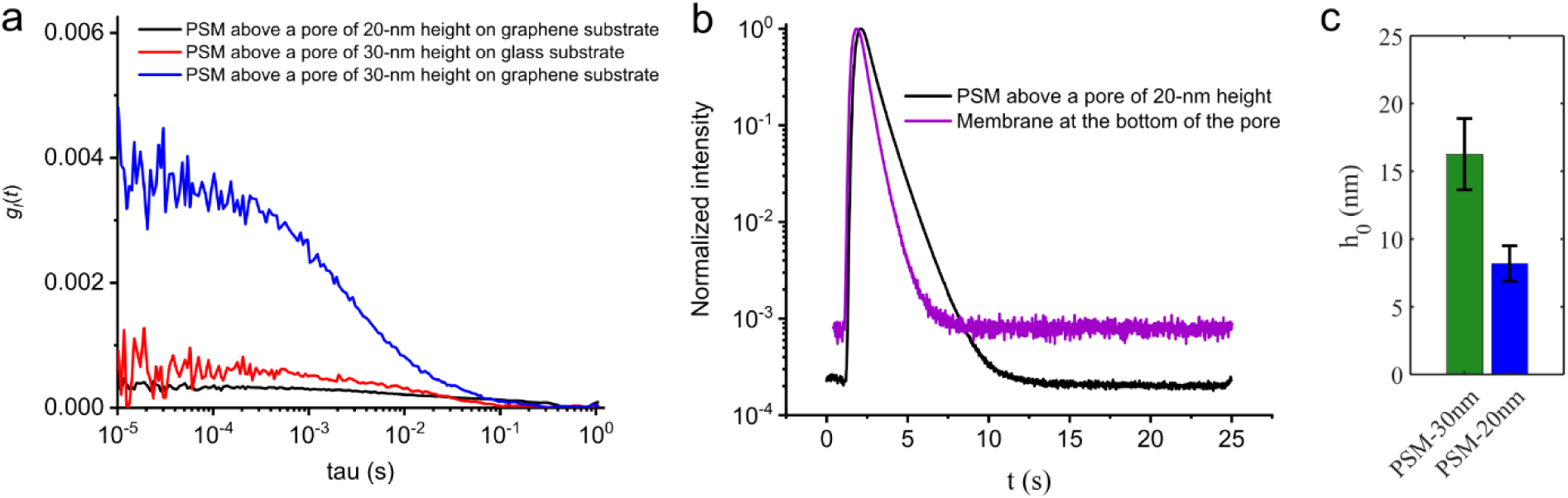
Intensity correlation and lifetime decay curves for PSM measurements. (a) Intensity correlation curves *gI*(*t*) measured at the center of the PSMs above different substrates. (b) Normalized fluorescence decay curves were for a PSM above a graphene-substrate with 20 nm deep pore, and for a membrane attached to the bottom of the pore. (c) Average height values (*h*0) for PSMs above prepared above pores with 30 nm and 20 nm depth.

**Supplementary Figure 7:**
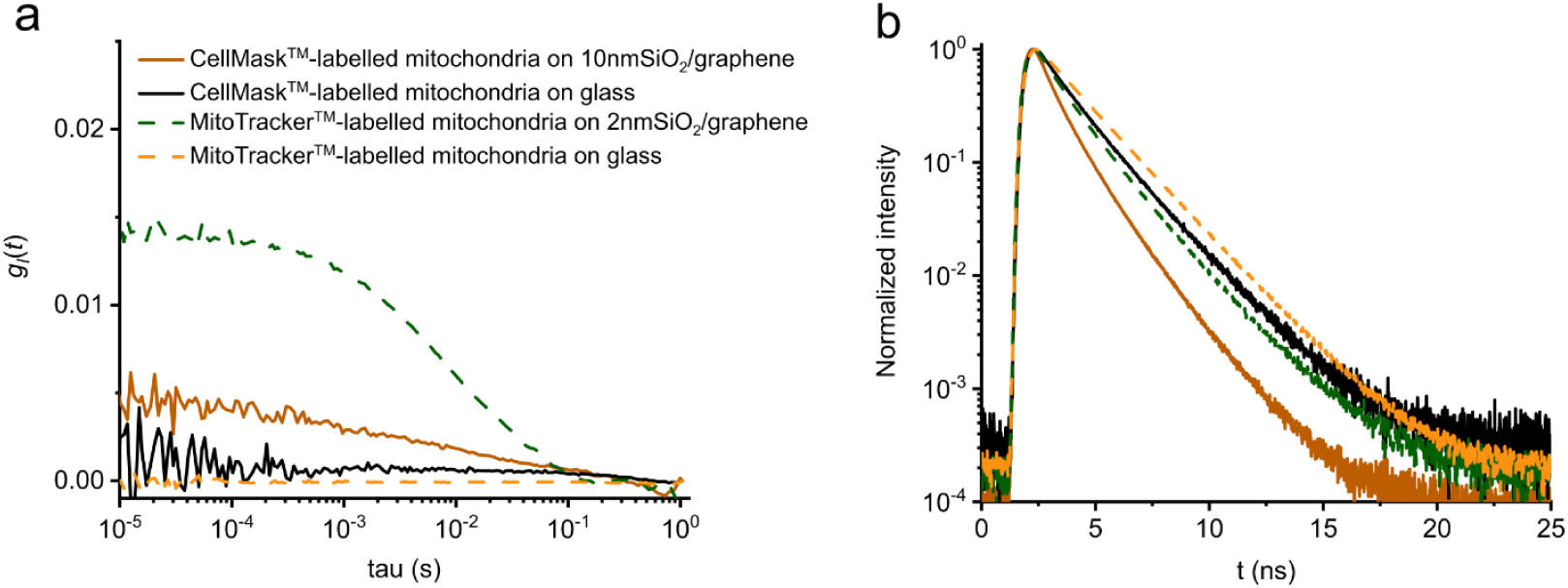
Intensity correlation and lifetime decay curves for mitochondria measurements. (a) Height correlation curves, *gh*(*t*), were measured for CellMask^TM^ and MitoTracker^TM^ labelled mitochondria in their active state, either a MIET substrate or on a glass coverslip. (b) Corresponding fluorescence lifetime decay curves.

**Supplementary Note 1: Validation of MIET lifetime and brightness curves**

To confirm the accuracy of the MIET lifetime and brightness curves, we conducted measurements on supported lipid bilayers (SLBs) with a well-known sample geometry. Specifically, we prepared SLBs of DOPC and 0.001% DPPE-Atto655 on four different substrates with silica spacer thickness values of 10 nm, 20 nm, 30 nm, and 50 nm. To account for the thickness of the SLBs and the hydration layer between the substrate and bottom leaflet of the SLBs, we calculated the lifetime and brightness curves assuming an average dye position of 2.5 nm above the surface (see Supplementary Figure 1a).

We took point measurements on these samples, with at least 10 measurements per sampling point. As shown in Supplementary Figure 1b and c, the measured lifetime values and count rates were found to be in excellent agreement with the theoretical calibration curves. These results confirm the accuracy of the MIET lifetime and brightness curves, and demonstrate their reliability for our experiments.

**Supplementary Note 2: Pore-spanning membrane (PSM) measurements**

To confirm that the pore-spanning membranes (PSMs) were free-standing, we employed two methods using negatively charged giant unilamellar vesicles (GUVs) containing DPhPC/DOPG/DPPE-Atto655 that were spanned over a positively charged pore substrate. The first method involved comparing the fluorescence correlation spectroscopy (FCS) curves obtained for PSMs above a glass substrate, the top membrane of GUVs (t-GUVs), and supported lipid bilayers (SLBs), all using the same lipid mixture. As shown in Supplementary Figure 5c, the FCS curves for t-GUVs and PSMs were almost identical, with faster diffusion rates than that of the SLBs.

The second method involved measuring the fluorescence lifetimes of the PSMs above graphene, see Supplementary Figure 6. The lifetime of the membrane touching the bottom of the pores was found to be very short, while the free-standing membrane had a longer lifetime. It should be noted that not all PSMs were suspended above the pore bottom, and only the data with long fluorescence lifetimes were used for further analysis.

Together, these results confirm that the PSMs were free-standing and not attached to the underlying substrate.

**Supplementary Note 3: Fluorescent labeling of mitochondrial membranes**

To label the outer and inner mitochondrial membranes (OMM and IMM), respectively, we used commercial dyes: CellMask™ Deep Red Plasma membrane Stain (Thermo Fisher) and MitoTracker™ Deep Red FM (Thermo Fisher). We employed the highest concentrations of the dyes recommended by the manufacturer (10 µg/mL for CellMask™ and 500 nM for MitoTracker™) to ensure optimal membrane labeling.

To confirm that these concentrations were sufficient to suppress fluorescence fluctuations due to lateral diffusion, we first measured the correlation function of the labeled mitochondria on glass substrates. As shown in Supplementary Figure 7a, the small amplitudes found on a glass substrate as compared to the larger amplitudes observed on a graphene substrate indicate that the lateral diffusion-related fluctuations originating from labeled OMM and IMM can be ignored. This confirms that the chosen dye concentrations are appropriate for our experiments and enables accurate interpretation of the results.

